# DDHD2 possesses both lipase and transacylase capacities that remodel triglyceride acyl chains

**DOI:** 10.1101/2025.09.22.677815

**Authors:** Lingshuang Wu, Yong Mi Choi, Mohyeddine Omrane, Jiyao Chai, Shujuan Gao, Abdou Rachid Thiam, Daniel Canals, Michael V. Airola

## Abstract

Hereditary spastic paraplegia subtype SPG54 is a genetic neurological disorder caused by mutations in the DDHD2 gene. Excessive lipid droplet accumulation is observed in the brains of SPG54 patients and DDHD2 knockout mice, consistent with DDHD2’s reported neutral lipase activity. Here, we find recombinant human DDHD2 preferentially hydrolyzes diacylglycerol (DAG) over phospholipids, with a slight preference for DAG over triacylglycerol (TAG). DDHD2 also exhibits transacylase activity, which enables transfer of acyl chains from triacylglycerols to diacylglycerols and monoacylglycerols to remodel the acyl chains of triglycerides. A predicted hydrophobic amphipathic helix on DDHD2 is essential for lipid droplet binding in vitro and in cells, and its lack reduces the enzymatic activity and triglyceride acyl chain remodeling. Adipose triglyceride lipase (ATGL), but not hormone sensitive lipase (HSL), also has transacylation activity and can remodel triglyceride acyl chains, but to a lesser extent than DDHD2. Taken together, this provides evidence that DDHD2 is a neutral lipid lipase and transacylase whose broad specificity enables triglyceride acyl-chain remodeling.

**SIGNIFICANCE STATEMENT:** Triglycerides (TAGs), the primary form of long-term energy storage, have acyl chain compositions crucial for diverse cellular processes. Lipases typically hydrolyze TAGs into free fatty acids. Here, we reveal a novel function for the neutral lipid lipase DDHD2: a transacylase activity. Instead of releasing fatty acids, DDHD2 transfers them between neutral lipids, altering TAG acyl chain composition. This transacylation requires the unique oil environment of lipid droplets (LDs), which excludes water from DDHD2’s lipolytic active site, favoring transacylation over hydrolysis. DDHD2’s lipase and transacylase activities enable TAG acyl-chain remodeling, demonstrating the possibility that a single enzyme can catalyze TAG cycling. This finding has implications for understanding lipid metabolism, LD dynamics, and specific motor neuron diseases implicating DDHD2.

## INTRODUCTION

Triglycerides (TAGs) are the primary form of stored energy in many organisms and are stored intracellularly within lipid droplets (LDs) [1–3]. Lipolysis is the catabolic process by which triglycerides are broken down into glycerol and free fatty acids [4]. The canonical lipases involved in lipolysis are adipose triglyceride lipase (ATGL) [5–8] and hormone sensitive lipase (HSL) [9, 10], which sequentially break down triglycerides. ATGL initiates the process by hydrolyzing triglycerides into diacylglycerol, followed by HSL which converts diacylglycerol into monoacylglycerol.

Triglyceride cycling is a metabolic process that involves the breakdown and resynthesis of triglycerides. This cycle is thought to be a “futile” cycle, because it does not alter overall triglyceride levels, but would consume energy through the consumption of acyl-CoAs during the resynthesis of triglycerides [11–13]. A recent study has traced triglyceride cycling using radiolabeled lipids in cells, but which enzymes may be involved in this pathway are unknown [14]. ATGL is recognized not only for its primary lipolytic activity but also for its capacity to perform transacylase functions. This dual functionality enables ATGL to catalyze the conversion of diacylglycerol (DAG) back to TAG [15–17]. Furthermore, ATGL’s bidirectional property extends to the metabolism of hydroxy fatty acids, allowing it to both synthesize and break down fatty acid hydroxy fatty acids (FAHFAs) [18, 19]. In contrast, the reports of enzymatic functions of HSL are limited to hydrolysis [20, 21]. This specialization allows HSL to process the lipid intermediates produced by ATGL, providing a downstream pathway for further lipid breakdown.

The enzyme DDHD2 (DDHD domain containing-2) has recently emerged as another lipase involved in the breakdown of neutral lipids. DDHD2 exhibits a broad tissue expression, with predominant levels in the brain, testis, and skeletal muscle [22]. This broad distribution suggests DDHD2’s potential involvement in diverse metabolic processes. Several studies have identified roles for DDHD2 in lipophagy [23], lipolysis [24–28], phospholipid metabolism [22, 28], vesicle trafficking [29–32], mitochondrial functions [33, 34] and synapse function [35].

Genetic studies have linked mutations in the DDHD2 gene with a neurodegenerative disorder, hereditary spastic paraplegia (SPG54), with patients characterized by a thin corpus callosum and lipid droplet accumulation in the brain [36–41]. In mouse models, DDHD2 knockout leads to lipid droplet accumulation in the brain [24, 25]. This led to the suggestion of DDHD2 functioning as a principal brain triglyceride lipase [24, 25]. However, other studies have suggested DDHD2 preferentially breaks down phospholipids or other neutral lipids like diacylglycerol (DAG) [24–27] and monoacylglycerol (MAG) [28], with a more recent study suggesting DDHD2 works in conjunction with ATGL and HSL to break down neutral lipids in various brain cell types [28]. The lack of a consensus substrate for DDHD2’s lipase activity remains an important unanswered question, which may vary by membrane environment (e.g. lipid droplets versus membrane bilayers).

In fact, the localization of DDHD2 to membrane structures has been shown to vary across different cell types. In HeLa cells, DDHD2 associates with the Golgi/ER-Golgi intermediate compartment [36]. In COS-7 cells, DDHD2 localizes to a prenuclear membrane structure that is adjacent to, but does not overlap with, markers for the ER or Golgi/ER compartments [25]. While in neuro-2a neuroblastoma cells, DDHD2 is enriched in the cytosol and partially on Golgi compartment [28]. Despite this variability in subcellular localization, DDHD2 has not been shown to co-localize with lipid droplets (LDs). This variability in DDHD2 subcellular localization suggests its dynamic nature and raises questions about the mechanisms that recruit DDHD2 to LDs.

Here, we examine the substrate specificity of human DDHD2 towards phospholipids and neutral lipids using in vitro assays and compared DDHD2’s enzymatic activity in different membrane environments with the canonical lipases ATGL and HSL. We found human DDHD2 is not only a lipase but can also function as a transacylase to synthesize triglycerides. We observe DDHD2 recruitment to lipid droplets in both U2OS and Huh-7 cells, and define a non-canonical amphipathic helix that is critical for both lipid droplet binding and transacylation activity. Lastly, we find that in vitro both DDHD2 and ATGL can remodel the acyl chains of triglycerides. Taken together, this study provides functional insight into how DDHD2 and ATGL proteins are lipases and transacylases and can potentially cycle triglycerides without ATP consumption.

## RESULTS

### DDHD2 preferentially hydrolyzes diacylglycerol over phospholipids

To examine the substrate specificity of human DDHD2, we purified His-tagged human DDHD2 from *E. coli* using Ni-NTA and size exclusion chromatography (Supplemental Fig. 1A) and first measured the enzymatic activity using fluorescent nitrobenzodiazole (NBD)-labeled lipids (Fig. 1A) solubilized in Triton X-100 mixed micelles. DDHD2 activity was compared between the lipids diacylglycerol (DAG), phosphatidylethanolamine (PE), phosphatidylcholine (PC), and phosphatidic acid (PA), as each lipid had previously been reported as a preferred substrate for either mouse [22] or rat DDHD2 [26]. The products were quantified using fluorescence-based high-performance liquid chromatography (HPLC), and all enzymatic activities were compared within the linear range of reaction time (Supplemental Fig. 1B) and protein concentration (Supplemental Fig. 1C).

**Figure 1.**
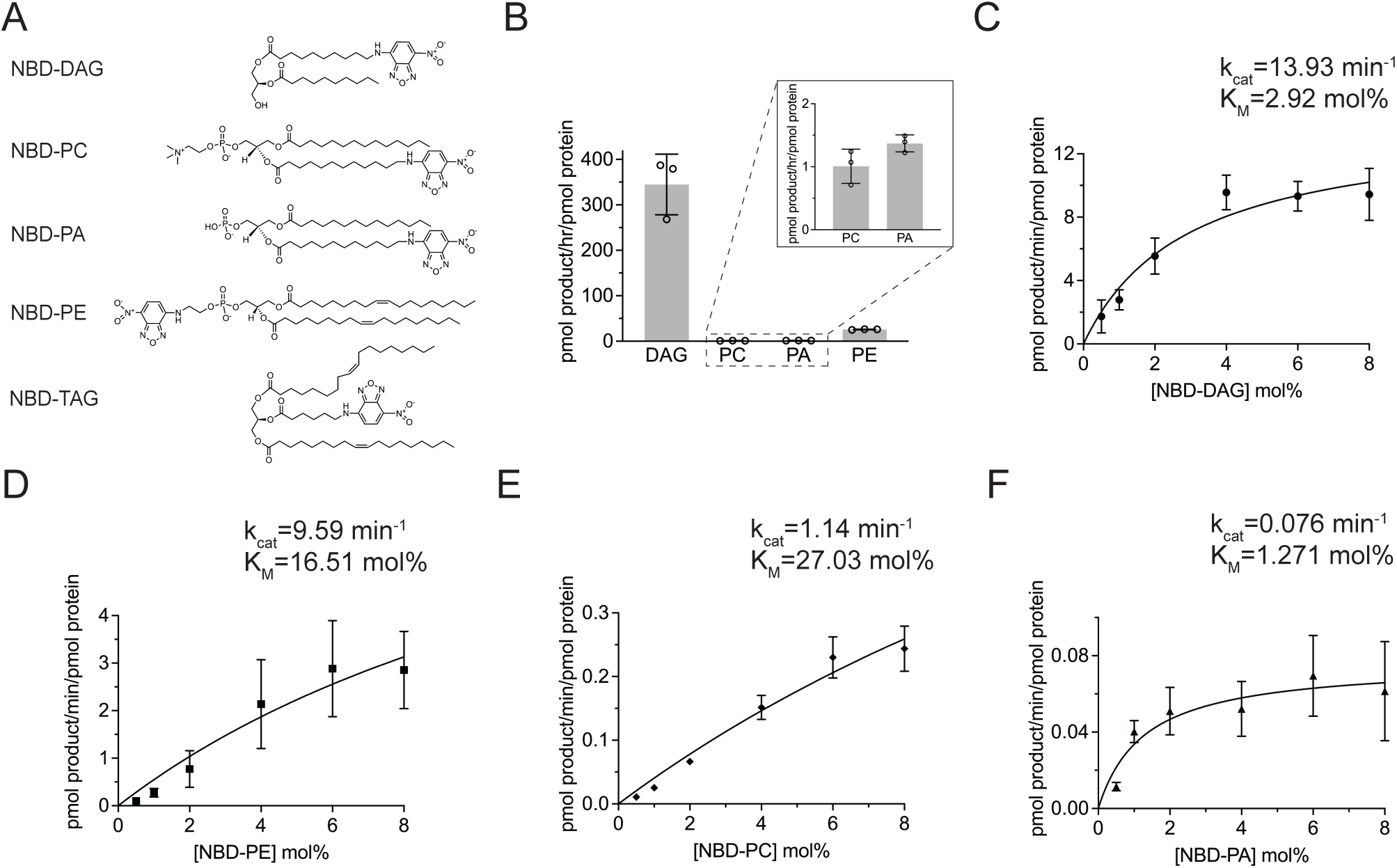
Human DDHD2 is a multifunctional lipase that preferentially hydrolyzes diacylglycerol over phospholipids. (A) Chemical structures of NBD-lipid substrates used in this study. DAG, diacylglycerol; PC, phosphatidylcholine; PA, phosphatidic acid; PE, phosphatidylethanolamine; TAG, triacylglycerol. (B) Quantification of DDHD2 activity against different lipids in Triton X-100 mixed micelles. Error bars represent standard deviation (n=3). (C-F) Effect of surface concentration (mol%) of DDHD2 activity towards (C) NBD-DAG, (D) NBD-PE, (E) NBD-PC, (F) NBD-PA in Triton X-100 mixed micelles. The surface concentration of NBD-lipids was varied by maintaining a constant bulk concentration of NBD-lipids (40 μM) and adjusting the amount of Triton X-100. Error bars represent SD (n=3).

DDHD2 showed a markedly higher hydrolytic activity towards DAG compared to PA or PC (Fig. 1B). Since the NBD group was on the sn-1 acyl chain of DAG, but on the sn-2 acyl chain of PA and PC (Fig. 1A), this raised the possibility that the higher activity towards DAG may be due to the acyl-chain position of the NBD group. Thus, we also tested activity against NBD-phosphatidylethanolamine (PE) with the NBD group as part of the head group and two unmodified oleate acyl chains (Fig. 1A). In comparison to PA and PC, DDHD2 displayed a ∼20-fold increase in activity against NBD-PE, which suggested the NBD acyl-linked moiety negatively affects DDHD2-mediated catalysis. However, despite this, DDHD2’s activity on NBD-PE was still significantly lower (nearly 13-fold) than NBD-DAG.

To further assess the substrate specificity of human DDHD2, steady-state kinetic parameters were determined (Fig. 1C-F). Among the tested substrates, NBD-DAG exhibited the highest catalytic efficiency (k_cat_/K_M_) which was ∼10-fold higher than NBD-PE, and ∼100-fold higher than both NBD-PC and NBD-PA. DDHD2’s preference for NBD-DAG as substrate derived from both a lower K_M_ of 2.9 mol%, compared to the estimated and extremely high K_M_’s of 16.5 mol% for NBD-PE and 27 mol% for NBD-PC, and DDHD2’s higher k_cat_ of 13.93 min^-1^ that far exceeded the k_cat_ of 0.076 min^-1^ for NBD-PA. Taken together, this suggests human DDHD2 preferentially hydrolyzes the acyl chain of DAG over phospholipids.

### Effect of membrane environment on DDHD2 catalysis

DDHD2 has been reported to act as a lipase at both intracellular membranes (e.g., the Golgi and plasma membrane), as well as on lipid droplets [25, 28, 42]. We thus sought to address whether the membrane environment influenced DDHD2 catalysis, with a particular focus on whether substrate preference was affected. Towards this goal, we assayed DDHD2 activity in liposomes and artificial lipid droplets (aLDs). Liposomes were generated using standard procedures [43]. In contrast, aLDs were generated using a previous protocol [44] with slight modifications that first solubilized the dried phospholipids in TAG oil by sonication, followed by the addition of aqueous buffer with two additional rounds of sonication (Supplemental Fig. 2A). Fluorescence microscopy of the aLDs showed low micrometer-sized aLDs that were stable for many hours (Supplemental Fig. 2B).

In liposomes, DDHD2 retained higher activity towards DAG as a substrate over all phospholipids tested (Fig. 2A). Still, it displayed only a 3-fold preference for NBD-DAG over NBD-PE, which could be affected by NBD-DAG being able to flip across the membrane bilayer [45], while NBD-PE cannot [46]. In contrast, in the lipid droplet system, the rate of NBD-DAG hydrolysis doubled, while the rate of NBD-PE and NBD-PC hydrolysis was severely diminished. This resulted in a ∼100-fold higher activity of DDHD2 for NBD-DAG over NBD-PE in lipid droplets. Taken together, this suggests that DDHD2 prefers to act on neutral lipids (e.g., DAG or TAG) versus the hydrolysis of the phospholipids in the lipid droplet monolayer.

**Figure 2.**
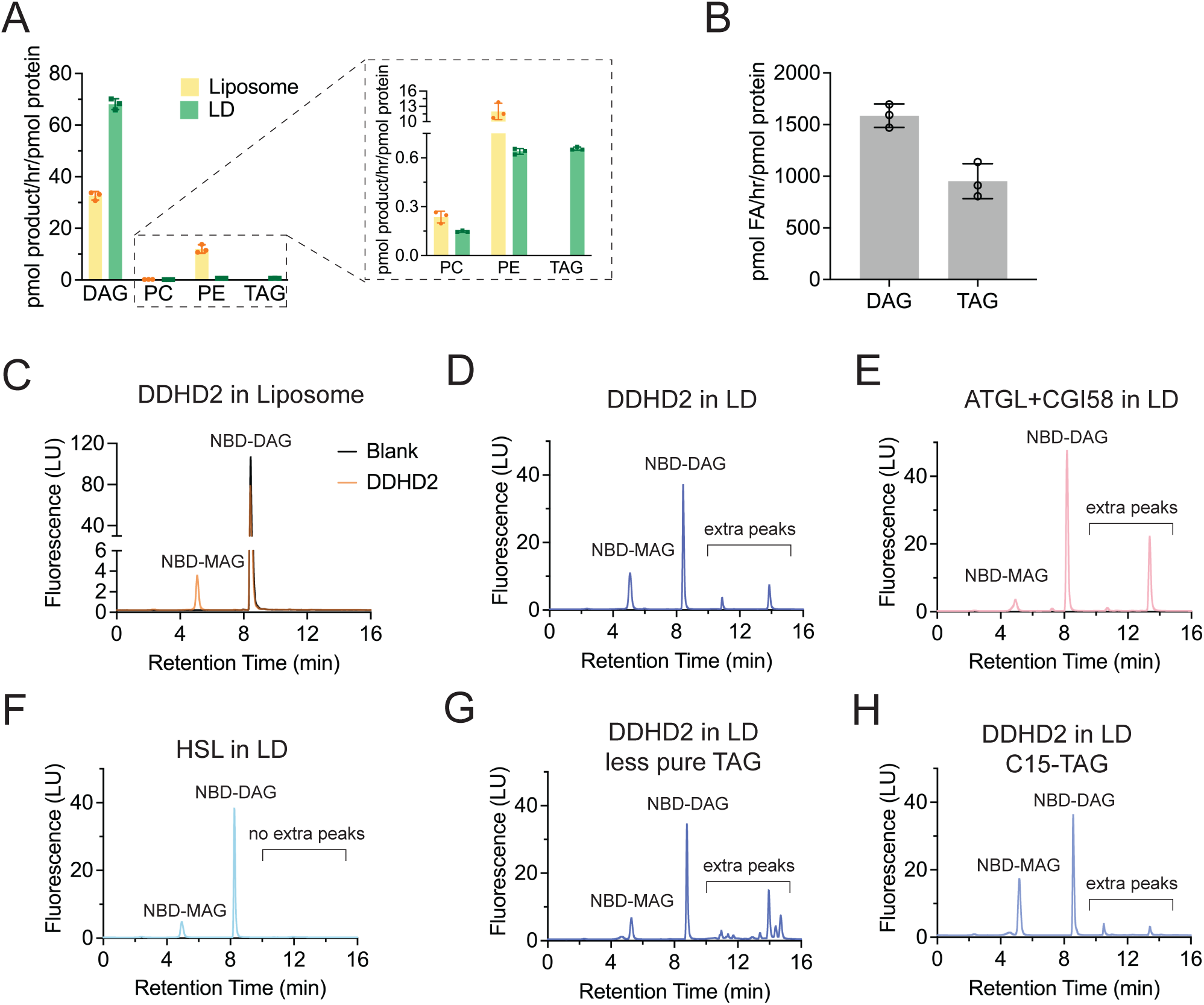
Effect of membrane environment on DDHD2 catalysis. (A) DDHD2 activity on NBD-labeled substrates including DAG, PC, and PE in liposome systems (bar and data points indicated in yellow) and DAG, PC, PE, TAG in artificial lipid droplet systems (bar and data points indicated in green). Data are presented as mean values ± SDs. n=3 independent experiments. (B) DDHD2 activity on the unlabeled substrates DAG (C18:1) and TAG (C18:1) quantified by measuring non-esterified fatty acid release. (C) Representative HPLC chromatogram of activity assay in liposomes. Blank yielded a single peak of the substrate NBD-DAG only (grey), while DDHD2 sample yielded an additional peak of the product NBD-MAG. (D) Representative HPLC chromatogram of activity assay in lipid droplets. Two additional more hydrophobic peaks were observed besides the NBD-DAG and NBD-MAG peaks. (E-H) Representative HPLC chromatograms of (E) ATGL with CGI58, (F) HSL, (G) DDHD2 with ∼50% pure TAG, and (H) DDHD2 with short chain C15-TAG activity assay on artificial lipid droplets using NBD-DAG as a fluorescent substrate.

We next analyzed DDHD2’s ability to hydrolyze NBD-TAG (Fig. 1A). In comparison to NBD-DAG, the activity of DDHD2 against NBD-TAG was low (Fig. 2A). As an alternative method, we also used the non-esterified fatty acid release measurement method to compare DDHD2’s preference between unlabeled DAG and TAG incorporated into Triton X-100 micelles. The results suggested a two-fold higher activity towards DAG over TAG (Fig. 2B).

### Characterization of DDHD2 transacylase and hydrolase activity

Using fluorescent NBD-labeled lipids as substrates allows for the direct monitoring of product formation and the potential of multiple products being generated simultaneously. In both mixed micelles and liposomes, DDHD2 hydrolyzed NBD-DAG to generate a single fluorescent product, NBD-monoacylglycerol (NBD-MAG) (Fig. 2C). However, in lipid droplets, DDHD2-mediated catalysis led to the emergence of not only NBD-MAG, but two additional fluorescent peaks (Fig. 2D). The longer retention times of these fluorescent peaks on the hydrophobic stationary column suggested DDHD2 could convert NBD-DAG (10:0-NBD/10:0) to at least two distinct, more hydrophobic products. We hypothesized this was due to transacylase activity of DDHD2 that would involve the transfer of an oleate acyl chain (C18:1) from the triolein TAG neutral lipid core used in our assay system to either NBD-DAG or NBD-MAG to form either NBD-TAG (10:0-NBD/10:0/18:1) or NBD-DAG (10:0-NBD/18:1).

To investigate this hypothesis, we compared the HPLC chromatograms of DDHD2 with two canonical lipases: hormone sensitive lipase (HSL) and adipose triglyceride lipase (ATGL). ATGL is known to exhibit transacylase activity and can transfer an acyl chain from TAG to DAG [15] or other hydroxyl-containing neutral lipids, including hydroxy fatty acids [18, 19]. HSL has not been demonstrated to exhibit transacylase activity, but HSL prefers to hydrolyze DAG as a substrate over TAG, similar to DDHD2. To carry out these experiments, recombinant human HSL was produced and purified from insect cells, and a truncated version of mouse ATGL [16, 47–49] and its co-activator CGI58 were purified from *E. coli*.

Using identical experimental conditions, we observed ATGL mediated catalysis, similar to DDHD2, resulted in the generation of two additional fluorescent peaks. However, the ratio of these peaks was notably different in comparison with DDHD2, with the major species generated by ATGL being the more hydrophobic fluorescent peak (Fig. 2E), which we suspected was NBD-TAG (10:0-NBD/10:0/18:1). In contrast, HSL-mediated catalysis generated a single product peak corresponding to NBD-MAG with no additional peaks observed (Fig. 2F). This suggested DDHD2, like ATGL, also exhibited transacylase activity and was capable of transferring acyl chains from TAG to DAG and potentially MAG.

To further validate our findings, we sought to alter the neutral lipid composition of the aLD oil core to confirm whether DDHD2 was transferring acyl chains from TAG or from the phospholipids used to generate the monolayer of the lipid droplets. Differentiating between these two possibilities was required, since under the above-described experimental setup, all lipids in both the neutral lipid core and the PC/PE phospholipid monolayer had oleate acyl chains with high purity (>95%). We reasoned that by varying the neutral lipid acyl chain composition, we may observe the generation of fluorescent products with variable retention times. Consistently, using a less pure (∼50%) triolein to form the neutral lipid core, but retaining the dioleoyl phospholipid monolayer, we observed the generation of multiple non-overlapping additional peaks (Fig. 2G). In addition, using a pure (>95%) but short chain (C15) TAG, we observed generation of two additional peaks that eluted at shorter retention times of 10.53 min and 13.46 min, compared to the retention times of additional peaks from pure triolein, which were 10.90 min and 13.86 min, respectively (Fig. 2H). We concluded that the putative transacylation activity of DDHD2 used the acyl chain of TAGs in the lipid droplet core, rather than the acyl chains from the phospholipid monolayer.

### DDHD2 is a transacylase that can cycle acyl chains of neutral lipids in lipid droplets

Thus far, we have demonstrated that DDHD2 can convert NBD-DAG to both NBD-MAG and additional fluorescent peaks, with this activity dependent on and requiring TAG. To directly characterize the chemical identity of the extra fluorescence peaks, we employed lipid chromatography-mass spectrometry (LC-MS). The chromatographic conditions, including the column type, solvents, and gradient profiles, were optimized and resulted in altered retention times and improved peak separation compared to our initial fluorescence-based HPLC protocol.

Qualitative analysis of NBD-labeled lipids was performed by analysis for their unique molecular fragmentation products using multiple reaction monitoring (MRM). The total ion chromatography revealed five individual peaks (Fig. 3A). Each lipid species was uniquely identified by the parent-to-daughter ion mass transition and corresponding LC retention time (Supplemental Table 1). This identified five chemically unique NBD lipids that corresponded to the original dodecanoyl NBD-DAG (10:0/10:0) substrate (Fig. 3C), the direct hydrolysis product NBD-MAG (10:0) (Fig. 3B), and three products derived from DDHD2 transacylation that involved transferring an oleate chain from triolein: NBD-DAG (10:0/18:1) (Fig. 3D), NBD-TAG (10:0/10:0/18:1) (Fig. 3E), and NBD-TAG (10:0/18:1/18:1) (Fig. 3F).

**Figure 3.**
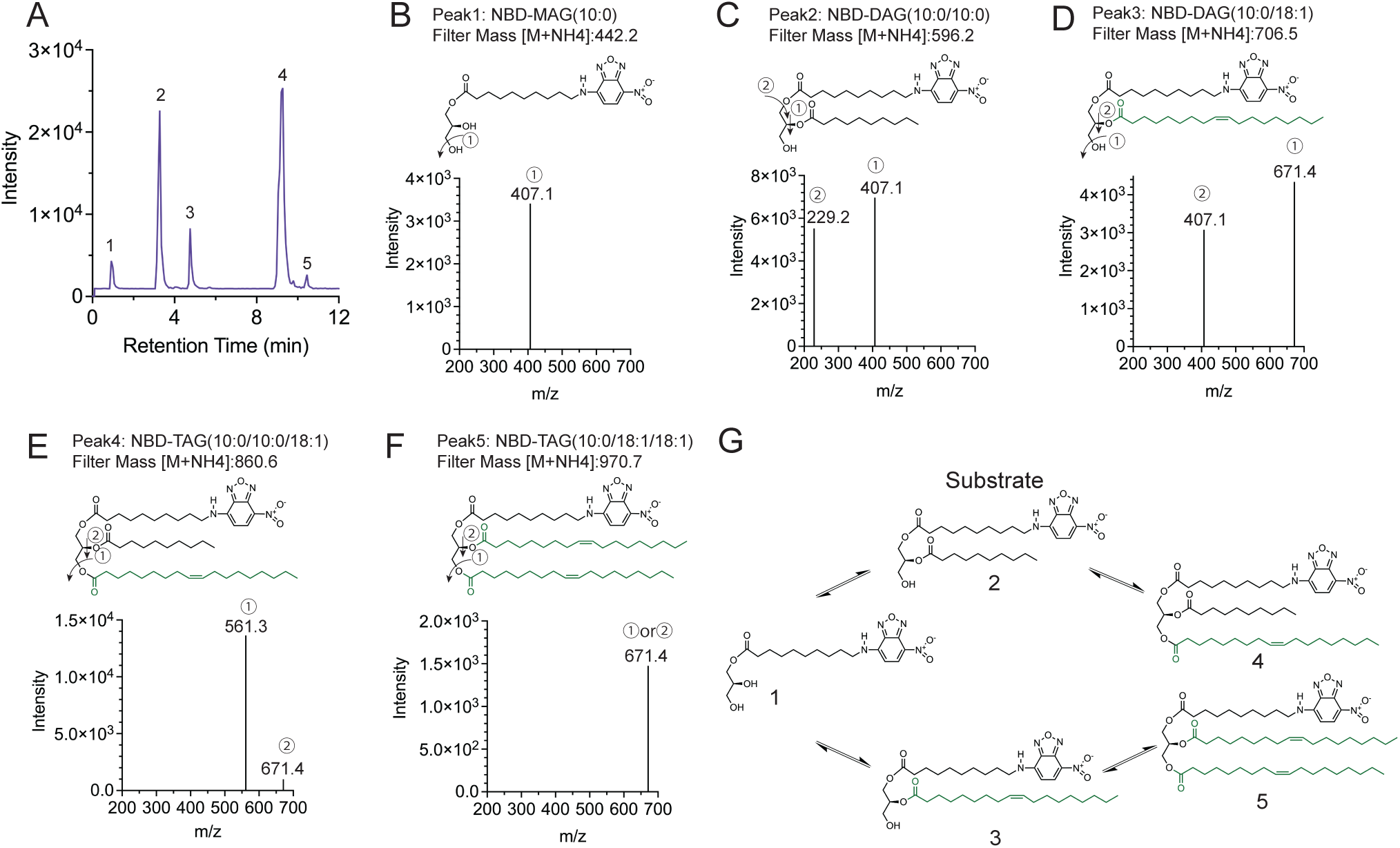
DDHD2 is a transacylase that can cycle triglyceride acyl chains. (A) Total ion chromatography for DDHD2 reaction using NBD-DAG as substrate. (B)-(F) Chemical structure and mass spectrum of (B) product NBD-MAG(10:0), (C) original substrate NBD-DAG(10:0/10:0), (D) product NBD-DAG(10:0/18:1), (E) NBD-TAG(10:0/10:0/18:1) and (F) NBD-TAG(10:0/18:1/18:1) filtered by mass [M+NH4+] indicated respectively, after collision energy 20 eV applied, molecule breaks down at position 1 or 2, detected mass indicated respectively. (G) Model for DDHD2 transacylation and hydrolysis activity on NBD-labeled substrates.

Together, these results provide evidence that DDHD2 exhibits dual enzymatic functions, acting not only as a hydrolase but also as a transacylase. Notably, the generation of NBD-TAG (10:0/18:1/18:1) required DDHD2 to both remove one 10:0 acyl chain by hydrolysis and add two 18:1 acyl chains through transacylation. While this is only demonstrated in vitro, this provides, to our knowledge, the first direct evidence of a lipase independently remodeling the acyl chains of triglycerides, without invoking the action of any acyl-CoA acyltransferases (Fig. 3G).

### Proposed catalytic mechanism(s) of DDHD2 in aqueous versus oil environments

Based on our results and structural predictions of DDHD2, we propose the following catalytic mechanism for the hydrolytic lipase and transacylase activities of DDHD2. Prior studies had identified DDHD2 as a serine hydrolase and AlphaFold [50] predicted DDHD2 to contain a catalytic triad consisting of serine, histidine, and aspartic acid residues, identical to the catalytic triad of canonical serine proteases whose mechanism is well described [51, 52]. Thus, during DDHD2 catalysis, we propose that Asp541 polarizes His681, which in turn deprotonates and activates Ser351 for nucleophilic attack of the carbonyl carbon within neutral lipid (e.g., DAG or TAG) ester linkages. This forms a tetrahedral intermediate (Fig. 4, Intermediate I) that subsequently breaks down to release a neutral lipid product (e.g., MAG or DAG) and forms an acyl-enzyme intermediate with the catalytic serine covalently bonded to a fatty acid.

**Figure 4.**
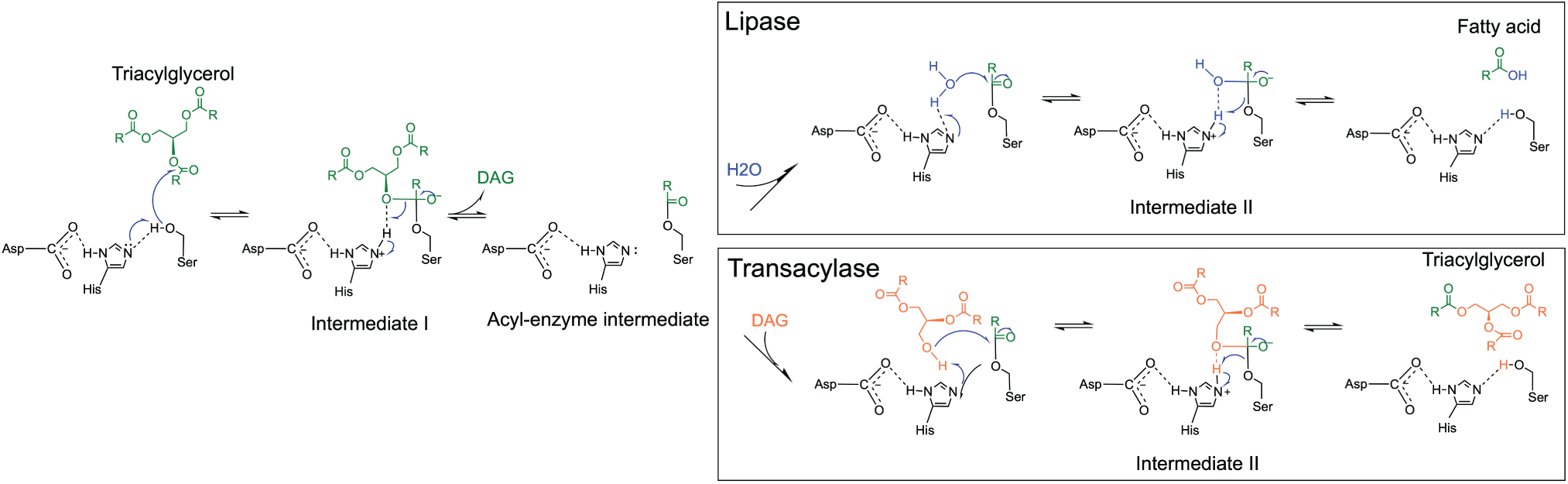
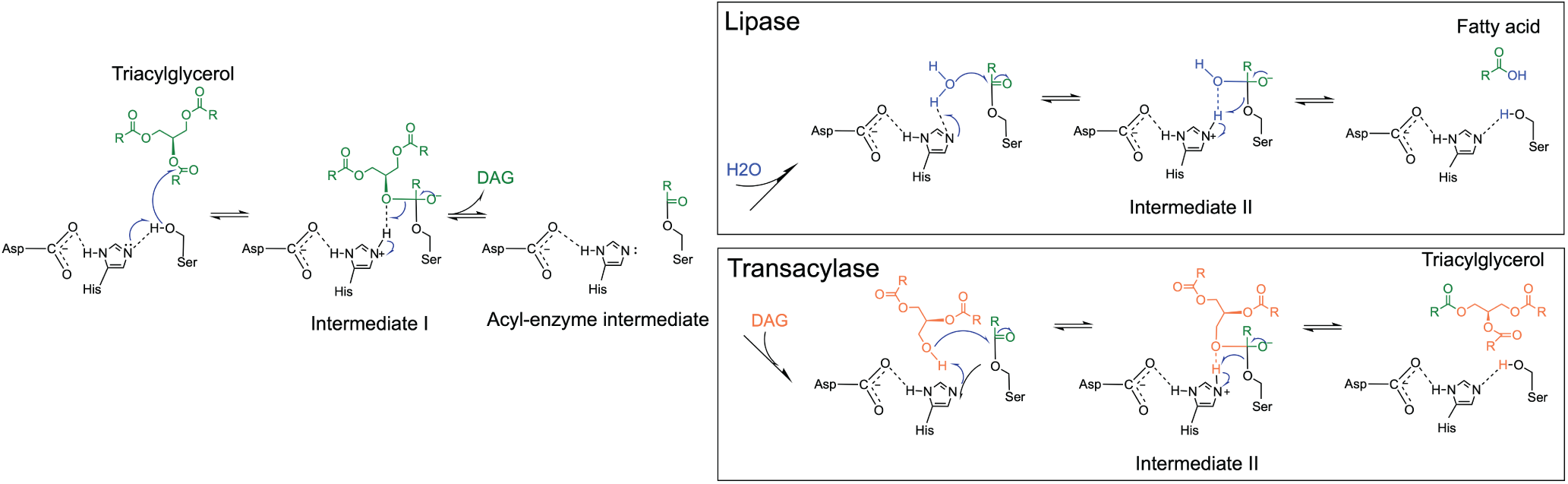
Proposed catalytic mechanism(s) of DDHD2 in aqueous versus oil environments. In an aqueous environment, DDHD2 lipase activity transfers an acyl chain of TAG to a water molecule (H2O) there by releasing a fatty acid (FA). In the oil membrane environment of lipid droplets, DDHD2 transacylation activity can occur due to exclusion of water, which would lead to transfer of an acyl chain of TAG to a DAG molecule, synthesizing new TAG.

Depending on the accessibility of the active site for water, the reaction can then proceed towards either hydrolysis or transacylation. In an aqueous environment, where water can enter the active site, hydrolysis is favored. The water molecule is deprotonated by His681 and activated for nucleophilic attack of the serine-acyl linkage, which proceeds to a second tetrahedral intermediate. Then it collapses, yielding a free fatty acid and regenerating the catalytic triad (Fig. 4, top right box).

Conversely, if water is excluded from the DDHD2 active site (e.g., if the DDHD2 active site is embedded in the oil environment of a lipid droplet), the reaction would stall at the acyl-enzyme intermediate until a hydroxyl group from an acceptor lipid (e.g., DAG) is deprotonated by His681 and activated for nucleophilic attack. This would lead to transacylation through formation and subsequent breakdown of a second tetrahedral intermediate (Fig. 4, bottom right box).

To verify the role of key residues in our proposed mechanism, we purified point mutations of Ser (S351A) (Supplemental Fig. 1A), His (H681A) and Asp (D541A) (Supplemental Fig. 3A). We found that all the point mutants were catalytically dead in both mixed micelles (Supplemental Fig. 3B) and aLDs (Supplemental Fig. 3C-F), supporting a critical role for these residues in catalysis.

### DDHD2 binds membranes via an alpha helix that is important for catalysis

In our proposed catalytic mechanism, the transacylase activity of DDHD2 depends on the exclusion of water from the active site during catalysis. We thus sought to test how altering the ability of DDHD2 to associate with membranes would affect the lipase and transacylase activities of DDHD2. We first sought to identify a region of DDHD2 involved in membrane association that was not directly involved in catalysis. Analysis of the AlphaFold structural prediction of DDHD2 identified an amphipathic helix present in a loop region of the catalytic domain (Fig. 5A,B). This helix was not a canonical amphipathic helix based on primary sequence (Fig. 5B) but was predicted to bend near a series of sequential branched-chain amino acids to form a continuous but curved amphipathic helix with classical opposing hydrophobic and hydrophilic surfaces (Fig. 5A).

**Figure 5.**
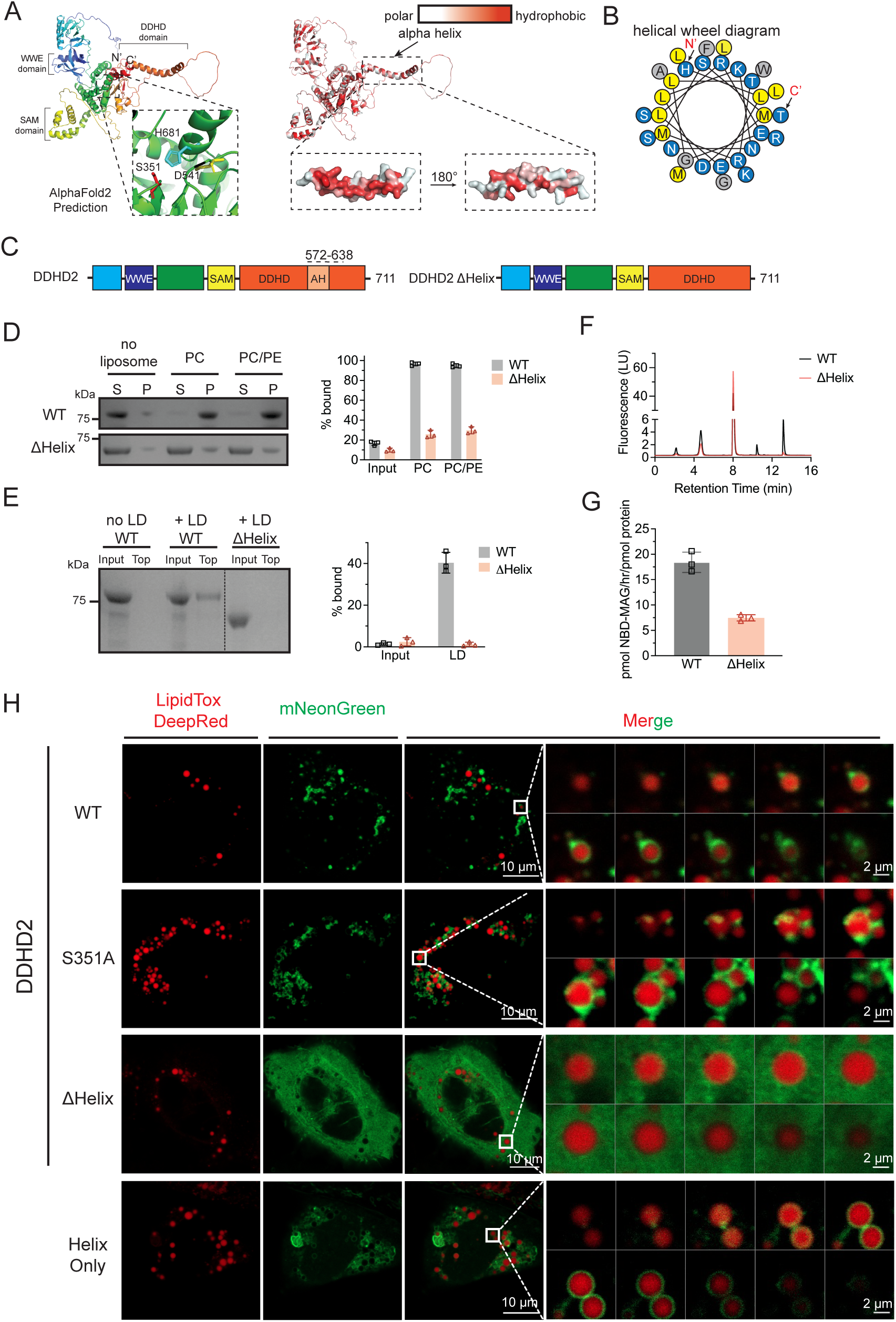
A non-canonical amphipathic alpha helix mediates DDHD2 membrane recruitment and catalytic activity. (A) Alphafold2 predicts human DDHD2 to form a typical serine hydrolase domain. The predicted DDHD2 structure is shown in cartoon form with rainbow coloring (left) from the N-terminus (blue) to C-terminus (red), or by hydrophobicity (right) with hydrophobic residues in red and hydrophilic residues in white. (B) Helical wheel diagram of the alpha helix. Hydrophobic residues are colored yellow or grey. Polar residues are colored blue. The position of the N- and C-termini are indicated. (C) Domain architecture of DDHD2 and DDHD2 ΔHelix. (D) SDS-page analysis (left) of liposome co-sedimentation assays reveal DDHD2 strongly associates with membranes, while deletion of the alpha helix in DDHD2 ΔHelix results in significantly less membrane association. S, supernatant; P, pellet; PC, phosphatidylcholine; PE, phosphatidylethanolamine. Quantification of liposome association (right) for DDHD2 and DDHD2 ΔHelix. Data are presented as mean values ± SDs. n=3 independent experiments. (E) SDS-PAGE analysis (left) of lipid droplet flotation assays reveals DDHD2 associates with lipid droplets, while deletion of the alpha helix in DDHD2 ΔHelix results in significantly less association. (F) Representative HPLC chromatography of DDHD2 ΔHelix (orange) compared to the DDHD2 wild type (grey) on lipid droplets. (G) Quantification of NBD-MAG produced by DDHD2 ΔHelix (orange) compared to the DDHD2 wild type (grey) on liposomes. (H) Left: Representative Airyscan images of U2OS cells stained by LipidTOX Deep Red (red) with overexpression of mNeonGreen-DDHD2 WT, mNeonGreen-DDHD2 S351A, mNeonGreen-DDHD2 ΔHelix and Helix-mNeonGreen only (green). Scale bars, 10 μm. Right: Zoom in of white box with Z-stack change from bottom to top. Scale bars, 1 μm.

Given that amphipathic helices mediate lipid droplet and membrane binding [53–56], we tested whether this non-canonical amphipathic helix was involved in DDHD2 membrane association. We purified DDHD2 lacking the amphipathic helix and an associated partially disordered loop (DDHD2 ΔΗelix) (Fig. 5C, Supplemental Fig. 1A) and first used liposome co-sedimentation to assess membrane binding. WT DDHD2 strongly associated with liposomes, displaying nearly 95% binding to PC or PC/PE containing liposomes. As predicted, deletion of the amphipathic helix in DDHD2 ΔΗelix severely reduced liposome binding (Fig. 5D). To verify that DDHD2 ΔΗelix also had reduced binding to aLD, we employed lipid droplet flotation assays. We found DDHD2 ΔΗelix did not bind to aLDs, in contrast to WT DDHD2, which demonstrated partial binding (Fig. 5E).

Having established that the ΔΗelix variant had reduced membrane association, we next tested the effects on catalysis. We found that DDHD2 ΔΗelix had a decrease in both hydrolysis and transacylation activity on lipid droplets compared to WT, with the transacylase activity nearly undetected (Fig. 5F). To directly compare hydrolysis activity, we also characterized activity using liposomes. We found the DDHD2 ΔΗelix activity was reduced by about 50% compared to WT (Fig. 5G).

To demonstrate the impaired membrane localization of the DDHD2 ΔHelix mutant, we generated mNeonGreen fusions of WT DDHD2, DDHD2 ΔHelix, the inactive S351A point mutant, and the helix alone. We assessed subcellular localization in two different cell lines that were treated with oleic acid (350 µM) to induce LDs in the cells: U2OS cells (Fig. 5H) and Huh7 cells (Supplemental Fig. 4).

We found WT DDHD2 partially localized around LDs (Fig. 5H, Supplemental Fig. 4). Notably, this is the first report observing DDHD2 localization to intracellular LDs. As noted by others [25, 28, 36], we also observed WT DDHD2 localized to other membranous structures at times. Deletion of the helix resulted in a complete loss of membrane and LD association (Fig. 5H), consistent with our in vitro binding data, while the helix alone localized directly on the surface of LDs (Fig. 5H). The S351A point mutant had enhanced localization to LDs in comparison to WT DDHD2 (Fig. 5H). Taken together, we concluded that the lack of the amphipathic helix in DDHD2 disrupts membrane binding and diminishes catalysis, with prominent effects on transacylase activity.

### DDHD2 can cycle triglycerides in vitro

Thus far, our data established that DDHD2 can function as both a lipase and a transacylase to synthesize NBD-TAGs and NBD-DAGs with mixed acyl chains. This implied that DDHD2 could cycle the acyl chain of triglycerides within LDs, and we sought to directly assess this using our in vitro system. To test this, we generated aLDs using approximately equimolar ratios of three different TAG species that each had three identical acyl chains that varied in length between them: 1,2,3-trioctanoyl glycerol (8:0/8:0/8:0), 1,2,3-trioleoyl glycerol (18:1/18:1/18:1), and 1,2,3-tri-12(Z)-heneicosanoyl glycerol (21:1/21:1/21:1). These aLDs were incubated with DDHD2 and the reaction products were analyzed using liquid chromatography-mass spectrometry (LC-MS) to identify and compare the resultant TAG species. As controls, we used a blank sample and the inactive S351A point mutant of DDHD2. We also included the DDHD2 ΔHelix, which had a more pronounced reduction in transacylation activity compared to hydrolysis activity.

After incubation with WT DDHD2, we identified seven new TAG species with acyl chains that were no longer uniform and were of variable length in the three different positions (Fig. 6A, B, Supplemental Table 2). Notably, these seven TAG species represented all seven possible TAG species combinations that could be generated from the initial homogeneous set of TAGs. We did not detect any new TAG species in the blank and S351A DDHD2 samples, indicating the observed triglyceride cycling was due to the enzymatic activity of DDHD2 and not from non-enzymatic acyl-chain migration. The DDHD2 ΔHelix could generate some new TAGs, but this was similarly profoundly reduced compared to WT DDHD2, consistent with the significant reduction in transacylation activity (Fig. 6B).

**Figure 6.**
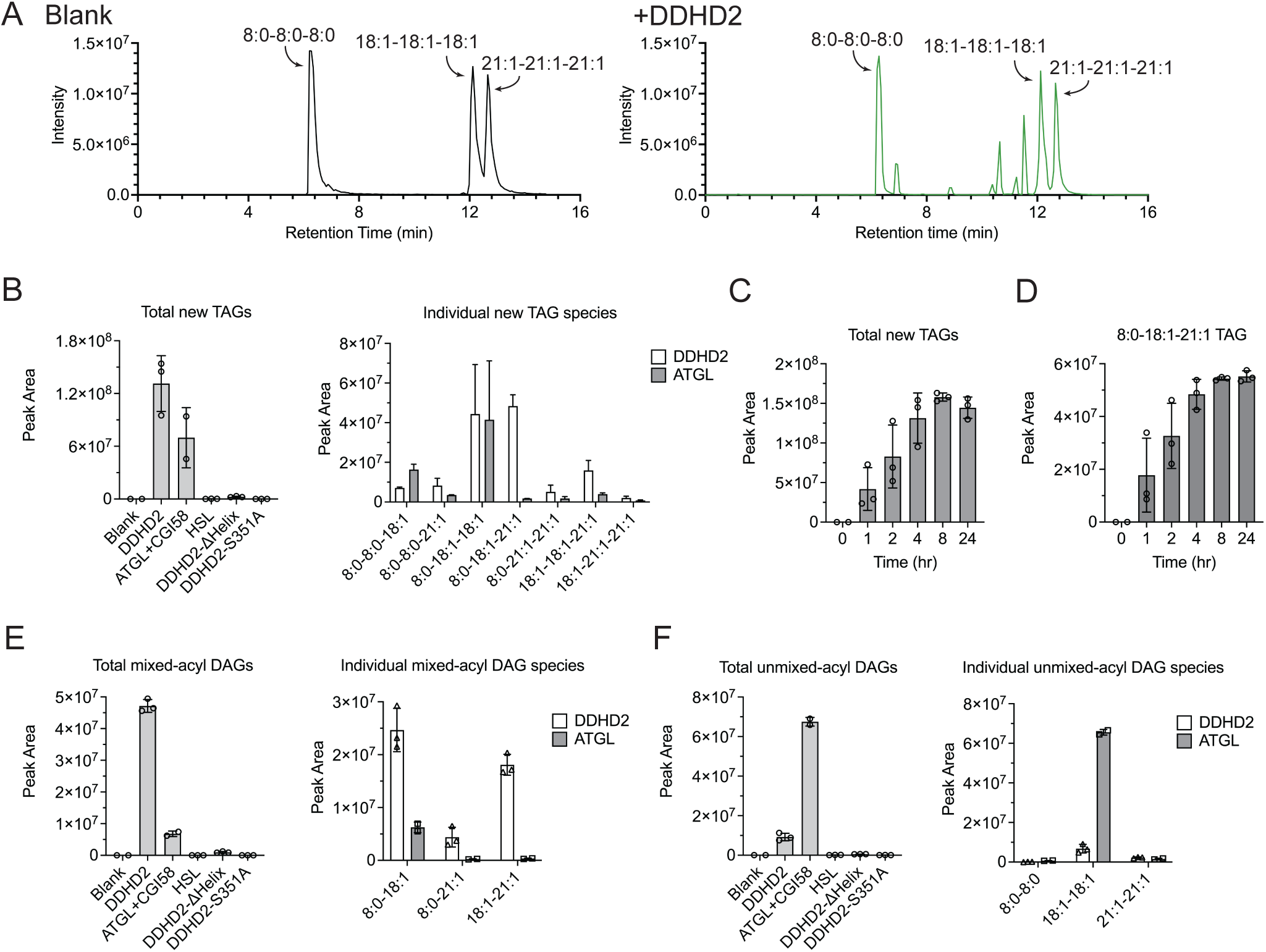
Acyl chain cycling of triglycerides by DDHD2 and ATGL. (A) Representative LC-MS total ion chromatogram of triglyceride cycling on lipid droplet made of 8:0/8:0/8:0, 18:1/18:1/18:1, and 21:1/21:1/21:1 TAGs. Left: blank, right: with DDHD2 incubation for 4 hrs. (B) Left: Total new TAGs generation in DDHD2(1μM), ATGL(10μM)+CGI58(10μM), HSL(1μM), DDHD2-ΔHelix(1μM), DDHD2-S351A(1μM). Right: Individual new TAG species in DDHD2 and ATGL in an order of increasing compound molecular weight, 8:0/8:0/18:1, 8:0/8:0/21:1, 8:0/18:1/18:1, 8:0/18:1/21:1, 8:0/21:1/21:1, 18:1/18:1/21:1, 18:1/21:1/21:1.(C) Total new TAGs and (D) 8:0/18:1/21:1 TAG on time scale from 1hr to 24hr in WT DDHD2 incubation. (E) Left: Total mixed-acyl DAGs generation by DDHD2 and ATGL+CGI58. Right: Individual mixed-acyl DAGs species including 8:0/18:1, 8:0/21:1, 18:1/21:1. (F) Left: Total unmixed-acyl DAGs generation by DDHD2 and ATGL+CGI58. Right: Individual unmixed-acyl DAGs species including 8:0/8:0, 18:1/18:1, 21:1/21:1.

A time course from 0 to 24 hours observed that after ∼4 hours, there were only relatively minor changes in the accumulation of new TAG species (Fig. 6C) and no significant changes in the 8:0-18:1-21:1 TAG species that had three acyl chains with unique lengths (Fig. 6D). We suspect this resulted from the reaction reaching equilibrium after ∼4 hours but cannot rule out that the DDHD2 enzyme may have lost enzyme activity over time.

We also observed the generation of new DAG species after incubation with DDHD2. We identified 6 distinct DAG species representing all 6 possible DAG species (Fig. 6E, F). Three of these DAG species could be the product of direct hydrolysis of TAG, with no changes in acyl-chain length from the TAG precursor (8:0-8:0 DAG, 18:1-18:1 DAG, and 21:1-21:1 DAG); while three of the species contained mixed acyl chain lengths (8:0-18:1 DAG, 8:0-21:1 DAG, and 18:1-21:1 DAG). Notably, we observed increased accumulation of the mixed acyl-chain length DAGs over the unmixed acyl-chain length DAGs after treatment with DDHD2 (Fig. 6E, F). This implies that DDHD2 also generated MAG that was subsequently transacylated to form the observed mixed acyl-chain length DAGs, similar to our results using 10:0-10:0 NBD-DAG, where we observed generation of 10:0-18:1 NBD-DAG.

### Comparison of triglyceride cycling by ATGL with DDHD2

ATGL has previously been characterized to exhibit both lipase and transacylation activities [5, 15], which was consistent with our observation of ATGL’s ability to convert NBD-DAG to NBD-TAG (Fig. 2E). We thus also compared the ability of ATGL to cycle triglycerides with DDHD2. HSL was also included as a control, given that HSL has not been reported to exhibit transacylation enzymatic activity.

An identical experimental setup generated lipid droplets with the three molecular TAG species: 8:0-8:0-8:0, 18:1-18:1-18:1, and 21:1-21:1-21:1 TAGs. Initial experiments with ATGL failed to detect a major accumulation of new TAG species. We thus repeated these experiments with a higher molar concentration of ATGL that was in 10-fold excess to the experiments used for DDHD2. ATGL’s co-activator CGI58 at a ratio of 1:1 with ATGL was also included to stimulate ATGL activity.

As observed for DDHD2, incubation with ATGL generated all seven possible new TAG species, while incubation with HSL gave rise to no appreciable new TAGs over the blank negative control. The total amount of new TAGs generated by ATGL was approximately half that observed for DDHD2. Notably, the number of individual TAG species that accumulated with ATGL was quite different from that of DDHD2. For example, DDHD2 and ATGL generated approximately equal amounts of 8:0-18:1-18:1 TAG, but DDHD2 generated ∼100-fold more of the completely mixed 8:0-18:1-21:1 TAG species than ATGL (Fig. 6B). We suspected that these differences may have arisen from the near equal specific activity of DDHD2 for hydrolysis of DAG and TAG, which would promote acyl-chain mixing. In contrast, ATGL’s preferred hydrolysis substrate is TAG. Consistent with this, we observed a higher increase in the generation of unmixed acyl-chain DAGs after ATGL treatment (Fig. 6F), with much more minor accumulation of the mixed-acyl chain DAGs (Fig. 6E). Overall, we concluded that ATGL, like DDHD2, is capable of remodeling the acyl chains of triglycerides. Still, the dual specificity of DDHD2 for both TAG and DAG enables DDHD2 to remodel triglycerides more efficiently than ATGL.

### DDHD2 has lipase and transacylase activity in cells

To determine whether the transacylase activity observed in vitro also occurs in cells, we developed a cell-based assay using overexpression of wild-type or catalytically inactive (S351A) DDHD2. Transfected U2OS (Fig. 7) and HEK293-T (Supplemental Fig. 5) cells were treated with 500 μM oleic acid for 3 hours to induce the formation of lipid droplets, after which DGAT1 and DGAT2 inhibitors were added to block endogenous TAG synthesis (Fig. 7A), and NBD-DAG was delivered to cells.

**Figure 7.**
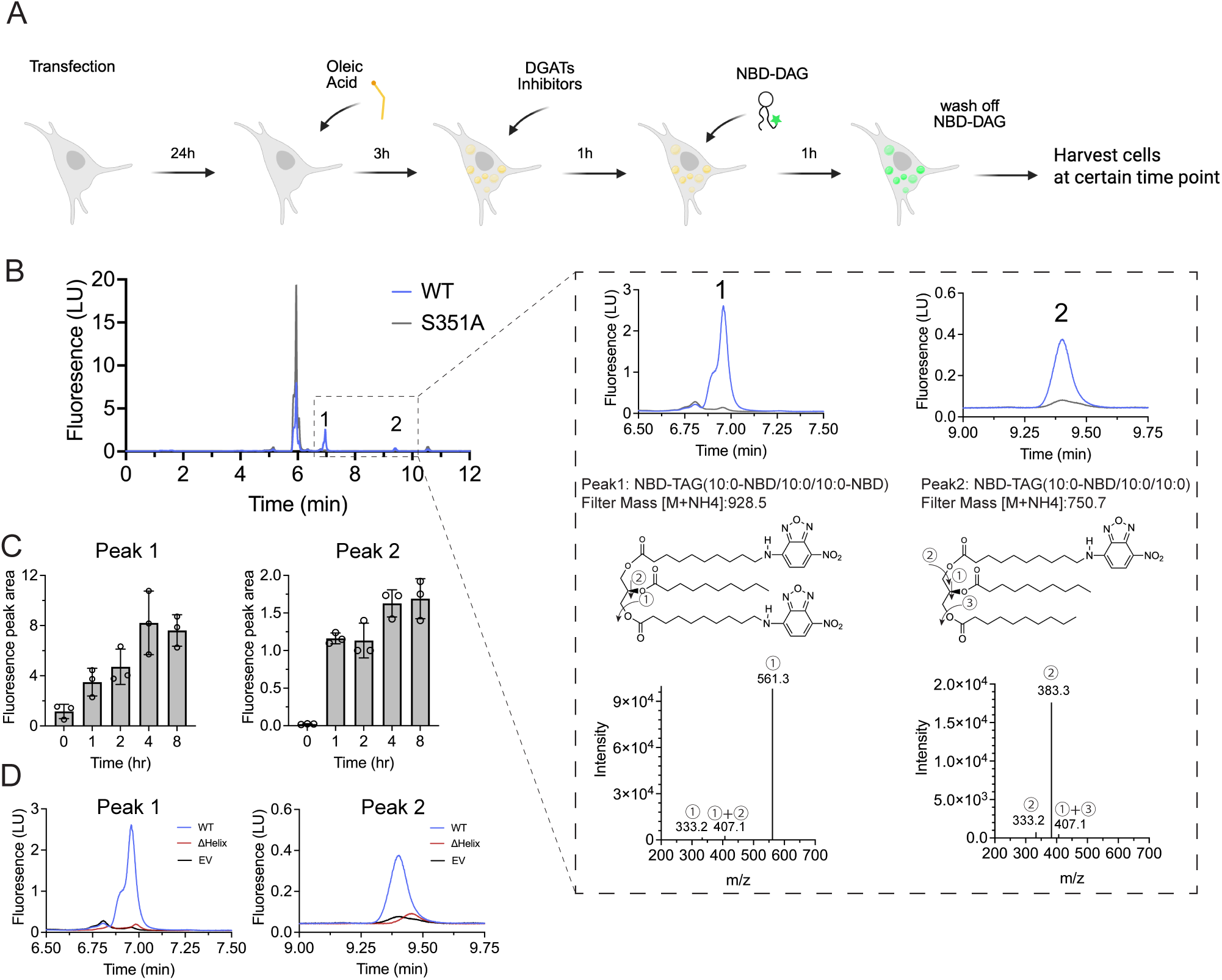
DDHD2 can generate new species of NBD-labeled triglycerides in cells. (A) Schematic graph of cell-based activity assay workflow. (B) Representative of HPLC fluorescence chromatography of cell-based activity assay of DDHD2 WT and inactive mutant S351A in U2OS cells. Two peaks generated only in DDHD2 WT are zoomed in. Peak 1 NBD-TAG (10:0-NBD/10:0/10:0-NBD) and peak 2 NBD-TAG (10:0-NBD/10:0/10:0) filtered by mass [M+NH4+] indicated respectively, after collision energy 20 eV applied, molecule breaks down at position 1, 2 or 3, detected mass indicated respectively. (C) Peak 1 NBD-TAG (10:0-NBD/10:0/10:0-NBD) and peak 2 NBD-TAG (10:0-NBD/10:0/10:0) on a time scale of 0 to 8 hours in DDHD2 WT incubation. (D) Zoom in peak 1 NBD-TAG (10:0-NBD/10:0/10:0-NBD) and peak 2 NBD-TAG (10:0-NBD/10:0/10:0) in DDHD2 WT, ΔHelix and empty vector.

In cells expressing DDHD2 WT, two new and distinct NBD-TAG species were observed by fluorescence. The chemical structures for these peaks were confirmed by LC-MS, with one peak being NBD-TAG (10:0-NBD/10:0/10:0-NBD) containing two NBD-10:0 acyl chains and one unlabeled 10:0 acyl chain, and the second species being NBD-TAG(10:0-NBD/10:0/10:0) with one NBD-labeled and two unlabeled 10-carbon acyl chains. The generation of these TAG species required DDHD2-catalyzed transacylation, where either an unlabeled 10-carbon or NBD-labeled acyl chain was transferred from one NBD-DAG molecule to another via transacylation in cells. Consistently, these peaks were absent in cells expressing the inactive S351A DDHD2 mutant (Fig. 7B, Supplemental Fig. 5A), as well as the DDHD2 ΔHelix variant and an empty vector containing only mNeonGreen (Fig. 7D, Supplemental Fig. 5B). A time course of 0 to 8 hours shows after ∼4 hours that an equilibrium may be reached as the levels of NBD-TAGs remained stable, and no longer increased (Fig. 7C). Taken together, these results demonstrate that DDHD2 is capable of catalyzing both the breakdown and remodeling of TAG and DAG within cells, thus defining DDHD2 as both an intracellular lipase and a transacylase.

## DISCUSSION

DDHD2 was originally characterized as a phospholipase that hydrolyzed the acyl chains of phospholipids [22, 28, 42]. However, after the observations that DDHD2 knockout mice accumulate LDs, there has been a growing body of evidence that DDHD2 preferentially hydrolyzes the neutral lipids TAG and DAG [24, 25, 28] rather than phospholipids. Here, by characterizing DDHD2’s lipase activity in different membrane environments, we provide more evidence that DDHD2 is a neutral lipid lipase that prefers to hydrolyze the neutral lipid DAG over phospholipids, with this preference further amplified by the membrane environment of lipid droplets. In line with prior work [26, 28], the minor, 2-fold preference for DAG over TAG suggests DDHD2 plays essential roles in both TAG and DAG breakdown, especially given that the concentration of TAG is much higher in lipid droplets than DAG. DDHD2 is thus a multi-functional neutral lipid lipase that combines the activities of both ATGL and HSL into a single enzyme capable of hydrolyzing both TAG and DAG at similar rates.

A major finding of this work is that DDHD2 exhibits transacylase activity. This enables DDHD2 the capability to synthesize and remodel the acyl chains of neutral lipids, e.g. via the transacylation of DAG to form TAG. This biochemical activity also raises the possibility that DDHD2 may synthesize an alternative, non-neutral lipid via transacylation, similar to the ability of ATGL to synthesize fatty acid hydroxy fatty acids (FAHFAs) [19] and PNPLA1’s synthesis of omega acyl-ceramide in skin tissues [57, 58]. A key question is whether the loss of transacylation contributes to the pathogenesis of HSP54, or whether it is simply the loss of DDHD2’s lipase ability that is causal in this neurodegenerative disease.

One key feature of the transacylation mechanism is that DDHD2 would need to bind to LDs to use TAG as an acyl donor and to access the oil environment of LDs to shift the equilibrium between hydrolysis and transacylation. Prior studies had not observed DDHD2 bound to LDs in cells, however, here we find DDHD2 can bind LDs and displays a LD-dependent transacylation activity within cells. We did observe a stronger co-localization of the inactive S351A DDHD2 mutant with LDs, and a reduction in the number of LDs after overexpression of WT DDHD2. These observations are consistent with the accumulation of TAG in HSP54 patients and the accumulation of LDs in DDHD2 KO mice. However, given that DDHD2 has been observed to bind multiple membranous cellular compartments, it remains a strong possibility that DDHD2 also has other non-overlapping functions in cellular lipid metabolism, which may include a role in lipophagy as recently reported [23].

At this point, there is sparse information about how DDHD2 lipase activity is regulated. However, DDHD2 regulation likely involves protein-protein interactions, similar to ATGL activation by CGI58 [6, 9] or HSL activation by perilipin1 [59, 60]. Alternatively, DDHD2 function could be attenuated by post-translational modifications, such as phosphorylation. If DDHD2’s activity is indeed regulated, this may have effects on DDHD2’s substrate specificity, preference for lipase or transacylation activity, and/or subcellular localization. Currently, there have been some reports of DDHD2 interacting with other proteins[23, 30], but this remains an active area of investigation.

## METHODS

### Materials

The following lipids were purchased from Avanti Polar Lipids: 1,2-dioleoyl-sn-glycero-3-phosphocholine (DOPC, catalog #850375), 1,2-dioleoyl-sn-glycero-3-phosphoethanolamine (DOPE, catalog #850725), 1,2-dioleoyl-sn-glycero-3-phosphoethanolamine-N-(7-nitro-2-1,3-benzoxadiazol-4-yl) (NBD-PE, catalog #810145), 1-Myristoyl-2-[12-[(7-nitro-2-1,3-benzoxadiazol-4-yl)amino]dodecanoyl]-sn-Glycero-3-Phosphocholine (NBD-PC, catalog #810123), 1-myristoyl-2-{12-[(7-nitro-2-1,3-benzoxadiazol-4-yl)amino]dodecanoyl}-sn-glycero-3-phosphate (NBD-PA, catalog #810172), N-[12-[(7-nitro-2-1,3-benzoxadiazol-4-yl)amino]dodecanoyl]-D-erythro-sphingosine (NBD-Ceramide, catalog #810211). 1-NBD-decanoyl-2-decanoyl-sn-Glycerol (NBD-DAG, catalog #9000341), 1,2,3-Tri-12(Z)-Heneicosanoyl Glycerol (catalog #26844), 1,2,3-Trioctanoyl Glycerol (catalog #26952), 1,2,3-trioleoyl glycerol (50% pure catalog #26871) was from Cayman Chemicals. 1,3-diolein, 2-NBD-X ester (NBD-TAG, catalog #6285) was from Setareh Biotech. Glyceryl trioleate (99% pure, catalog #T7140) was purchased from Millipore Sigma. Q5 Site-Directed Mutagenesis Kit and NEBuilder® HiFi DNA Assembly Master Mix were purchased from New England Biolabs. Nickel-NTA Agarose resin was a product of GoldBio.

### Protein expression and purification

Full length recombinant human DDHD2, the S351A, D541A and H681A point mutants, and the human DDHD2 ΔHelix were produced and purified from *E. coli*. Mouse ATGL 1-319 and full length human CGI58 were also produced and purified from *E. coli*. Recombinant human HSL was produced and purified from Sf9 cells using baculovirus. Full details on the methods for overexpression and protein purification are provided in the supplemental information.

### Substrate preparation

Mixed Micelles - NBD-labeled lipid substrates in chloroform were dried under nitrogen gas. Triton X-100 was used to generate mixed micelles in buffer containing 50 mM HEPES pH 8, 50 mM NaCl. The mixture was sonicated in a water bath for 30 min. The substrate mixed micelle mixtures contained 1.25 mM Triton X-100 and 25 μM NBD-labeled lipid substrates, which were mixed 50:50 with protein solution. The final concentration of NBD-lipids in assays was 12.5 μM, unless otherwise noted. For Michaelis-Menten kinetic experiments, the final concentration of NBD-labeled lipids was held constant at 20 μM and the amount of Triton X-100 was varied. This varied the surface concentration of NBD-lipids in each Triton X-100 mixed micelle, which we express as a molar percentage (mol% NBD-lipid).

Liposomes - Liposomes were generated using standard procedures [43]. Large unilamellar vesicle (LUV) liposomes were prepared for the liposome system. The lipids DOPC, DOPE, and NBD-labeled lipid substrates in chloroform were mixed in a molar ratio of 80:20:2 and dried under nitrogen gas. The lipids were resuspended in buffer containing 50 mM HEPES pH 8, 50 mM NaCl and LUVs were generated by 7 freeze-thaw cycles followed by a water-bath sonication. The final substrate liposome mixture contains 1 mM DOPC, 0.25 mM DOPE, and 25 μM NBD-labeled lipid substrates.

Lipid droplets - aLDs were generated using a previous protocol [44] with slight modifications. A calculated amount of DOPC and DOPE was mixed with NBD-labeled substrate in a molar ratio of 80:20:2. The mixture was dried under nitrogen gas to achieve a final concentration of 1 mM DOPC, 0.25 mM DOPE, and 25 μM NBD-substrate. 10 μL Glyceryl trioleate (Triolein) in neat oil state was added to the dried lipids and water bath sonicated until all lipids were emersed in the oil. 500 μL of a buffer solution, which contained 50 mM HEPES at pH 8 and 50 mM NaCl, was added. This was followed by sonication in a water bath, intermittent vortex mixing for 20 minutes, and tip sonication for 10 minutes at 35% amplitude with intervals of 2 seconds on and 2 seconds off. To visualize under fluorescence microscope, aLDs were prepared by mixing DOPC, DOPE, Rhodamine-PE and NBD-DAG in a molar ratio of 80:19:1:2 at the same final concentration as above. The aLD solution was diluted 100-fold, added onto BSA-blocked glass slides, and imaged using a ZEISS LSM 980 confocal microscope equipped with Airyscan.

### In vitro enzymatic activity assay for liquid chromatography

The prepared substrate (micelle, liposome or lipid droplet) and protein were mixed at 1:1 volume ratio and incubated for 1 h at 37°C. The reaction was terminated by addition of a methanol:chloroform (1:1) solution and vortexing vigorously to mix. The mixture was centrifuged at 3000 x g for 5 min, the lower organic phase was extracted using a glass Hamilton syringe and transferred to a new glass tube. After the organic phase was dried under nitrogen gas for 30 min and resuspended in HPLC Buffer B containing 100% methanol, 1 mM ammonium formate and 0.2% formic acid (v/v). The solution was centrifuged at 3000 x g for 5 min and the supernatant was transferred to an HPLC vial with inserts.

### In vitro enzymatic colorimetric assay

Unlabeled neutral lipid (DAG or TAG) was emulsified into Triton X-100 micelles by several rounds of water bath and tip sonication. The substrate and protein were mixed at 1:1 volume ratio and incubated at 37°C. The activity assay buffer used was comprised of 150 mM Tris-HCl pH 8, 500 mM NaCl, and 1% Triton X-100. The fatty acid release was measured following the manufacture protocol (FUJIFILM, Wako HR series NEFA-HR(2)). Purified aliquots of the DDHD2 protein were used to determine the linearity of the reaction over time, which was linear up to 90 minutes of total reaction time. The protein concentration linearity of DDHD2 was also determined, ranging from 0.1 μM to 0.4 μM. The amount of non-esterified fatty acids release in each condition was quantified using a standard curve of oleic acid determined using the activity assay buffer.

### Liposome sedimentation assay

40 μL of LUV liposomes in 50 mM Tris-HCl pH 8, 150 mM NaCl, 1 mM TCEP buffer were mixed with 20 μL of proteins to give a final concentration of 0.67 mM liposomes and 4 μM protein. Reaction mixtures were incubated for 40 mins and centrifuged at 100,000 x g at 4°C for 1h. The supernatant fraction was carefully collected, and the pellet was resuspended in the same volume of buffer, then analyzed by SDS-PAGE. Binding assays were performed more than three times, and the gel bands intensity were quantified using ImageJ.

### Lipid droplet floatation assay

160 μL of lipid droplets in 50 mM HEPES pH 8, 50 mM NaCl buffer were mixed with 40 μL of proteins to make a final molar ratio of 1 protein to 1000 phospholipid. After 1h incubation on ice, an 50 μL input fraction was removed. The remaining reaction was mixed with 1:1 volume ratio with 80% Nycodenz (catalog # YD07039) and loaded into the bottom of a Beckman centrifuge tube (catalog # Z71229SCA). 300 μL of 30% Nycodenz was slowly layered on the top of the bottom fraction. Another 100 μL buffer was then layered on the top of the fraction then centrifuged at 197,300 x g in a Beckmann SW55 swinging bucket rotor for 4h. Top, middle, bottom layers were carefully taken out and equal percentage with input fraction are separately mixed with LDS loading dye and loaded on SDS-PAGE gel. The gel bands intensity was quantified using ImageJ.

### Fluorescence microscopy

The following plasmids were used for fluorescence microscope experiments.

mNeonGreen-DDHD2 WT,
mNeonGreen-DDHD2 S351A,
mNeonGreen-DDHD2 ΔHelix (aa 572-638),
DDHD2-Helix-mNeongreen (aa 567-638 of DDHD2),
mNeongreen-ATGL.

The above constructs were prepared with mNeonGreen-mTurquoise2 vector (mNeonGreen-mTurquoise2 was a gift from Dorus Gadella (Addgene plasmid # 98886; http://n2t.net/addgene:98886; RRID:Addgene_98886) [61] by Gibson Assembly.

Bovin serum albumin-conjugated oleic acid (OA-BSA) was prepared as described [62]. Briefly, oleic acid was diluted with 100% ethanol to 500mM and 10% (w/v) BSA was prepared with fatty acid free BSA in DMEM. Oleic acid was then diluted with 10% BSA to a final concentration of 5 mM.

U2OS cells were grown at 37°C in 5% CO2 in Dulbecco’s modified Eagle’s Medium (DMEM) (Corning) supplemented with 10% fetal bovine serum (FBS) and 1% penicillin/streptomycin. Cells were used for experiments between passage 5 to 25. Cells were seeded at 60% confluence in glass bottom dishes (Cellvis) in DMEM with 10% FBS and cultured overnight before transfection. Cells were transfected with corresponding plasmids using Lipofectamine 3000 (Invitrogen) based on manufacturer recommendations. Cells were cultured overnight in DMEM with 10% FBS. Cells treated with oleic acid were supplied with 350 μM OA-BSA 3h after transfection. Lipid droplet was stained with HCS LipidTOX Deep Red (Invitrogen) at 1:5000 dilution in PBS for 15 minutes at 37 °C. Cells were then fixed with 4% paraformaldehyde (PFA) in phosphate-buffered saline (PBS) for 15 minutes at room temperature, followed by washing with 1mL PBS for three times. The cells were either imaged immediately or kept at 4 °C before imaging. Cells were imaged with a Zeiss LSM 980 Airyscan 2 NLO Two-Photon Confocal Microscope at Stony Brook Central Microscopy Imaging Center.

Huh7 cells were maintained in High Glucose / stabilized Glutamine and Sodium Pyruvate Dulbecco’s modified Eagle’s Medium (DMEM) (Gibco) supplemented with 10% heat-inactivated fetal bovine serum and 1% penicillin/streptomycin (GibcoBRL) (complete media) at 37°C and 5% CO2. Huh7 cells were seeded at 60–70% confluence in MatTek 3.5mm coverslip bottom dishes and incubated in DMEM complete media overnight before transfection. Cells were transfected corresponding plasmids using X-tremeGENE™ 9 based on the manufacturer recommendations. Cells were incubated overnight in DMEM complete media supplemented with Oleic Acid (OA)/Bovin Serum Albumin (BSA) complex at corresponding concentration of 200 µM of OA in order to induce lipid droplet accumulation then cells were incubated in Earle’s Balanced Salt Solution (EBSS)(Gibco) for further 24h. HCS LipidTOX™ Neutral Lipid Stains (Invitrogen) was used to stain lipid droplets (1/2000) for 30 minutes at 37°C before imaging. Live cells were imaged with a ZEISS LSM 9 with Airyscan.

### Cell-based activity assay

U2OS cells were plated and transfected as described above. HEK293-T cells were transfected with corresponding plasmids using X-tremeGENE 9 DNA Transfection Reagent (Sigma) based on manufacturer recommendations. Cells were cultured in DMEM with 10% FBS for 24 h. 500 μM OA-BSA was used to treat cells for 3h to form lipid droplets. OA-BSA was removed by change media into DMEM with 20μM DGAT1 inhibitor (PF-04620110, Cayman) and 20μM DGAT2 inhibitor (PF-06424439, Cayman). After 1 h, 20μM NBD-DAG (Cayman) was added to the cell culture. After 1 h, NBD-DAG was removed by change media into DMEM with DGATs inhibitors. Cells were then washed with ice-cold PBS three times and collected by cell scraper with the addition of 500 μL of extraction mix, Isopropanol: ethyl acetate: H2O (6:3:1) with 250 pmols internal standard (d5-Triolein, Cayman) in glass tube. The solution was dried under N2 gas until complete dry, then methanol:chloroform (1:1) solution was added and vortexed vigorously to mix. The mixture was centrifuged at 3000 x g for 5 min, the lower organic phase was extracted using a glass Hamilton syringe and transferred to a new glass tube. After the organic phase was dried under nitrogen gas for 30 min and resuspended in HPLC Isopropanol mobile phase B (1% H2O / 9% acetonitrile / 90% isopropanol (v/v/v),10 mM ammonium formate). The sample were further injected into HPLC with fluorescence detector or LC-MS for further analyses.

### HPLC fluorescence analyses

The samples from in vitro enzymatic activity assay were injected in an Agilent HPLC system with Spectra 3 μm C8SR column (Catalog # S-3C8SR-FJ). Injection volume: 10 μL; Flow: 0.5 mL/min; Mobile phase A: Fisher Water Optima LC/MS, 1 mM ammonium formate, 0.2% formic acid (v/v); Mobile phase B: Fisher Methanol Optima, 1 mM ammonium formate, 0.2% formic acid (v/v); Gradient: 0-1 min 50% to 80% Buffer B, 1-8 min 80% to 98% Buffer B, 8-11 min 98% Buffer B, then 11-16 min 80% Buffer B. The fluorescence detector was set to scan excitation wavelength 470 nm and emission wavelength 530 nm according to the properties of the NBD fluorophore.

The samples from cell-based activity assay were injected in an Agilent HPLC system with Poroshall 120, EC-C18, 2.7 μm column (Catalog # 695975-302). Injection volume: 10 μL; Flow: 0.4 mL/min; Mobile phase A: 50% H2O / 30% acetonitrile / 20% isopropanol (v/v/v),10 mM ammonium formate; Mobile phase B: 1% H2O / 9% acetonitrile / 90% isopropanol (v/v/v),10 mM ammonium formate; Start with 90% solvent A, 10% solvent B; Gradient: 0-2.7 min 10% to 45% solvent B, 2.7-2.8 min 45% to 53% solvent B, 2.8-9 min 53% to 65% solvent B, 9-9.1 min 65% to 89% solvent B, 9.1-11 min 89% to 92% solvent B, 11-13.9 min 92% to 100% solvent B, 13.9-14.9 min 100% to 10% solvent B. The fluorescence detector was set to scan excitation wavelength 470 nm and emission wavelength 530 nm according to the properties of the NBD fluorophore.

### LC-MS analyses

The samples from in vitro enzymatic activity assay were injected in an Agilent 1290 series HPLC system with ZORBAX RRHD Eclipse Plus C18 1.8μm column (Catalog # 959758-902) Injection volume: 3 μL. The samples from cell-based activity assay were injected in Poroshall 120, EC-C18, 2.7 μm column (Catalog # 695975-302). Injection volume: 10 μL. Flow: 0.4 mL/min; Mobile phase A: 50% H2O / 30% acetonitrile / 20% isopropanol (v/v/v),10 mM ammonium formate; Mobile phase B: 1% H2O / 9% acetonitrile / 90% isopropanol (v/v/v),10 mM ammonium formate; Start with 90% solvent A, 10% solvent B; Gradient: 0-2.7 min 10% to 45% solvent B, 2.7-2.8 min 45% to 53% solvent B, 2.8-9 min 53% to 65% solvent B, 9-9.1 min 65% to 89% solvent B, 9.1-11 min 89% to 92% solvent B, 11-13.9 min 92% to 100% solvent B, 13.9-14.9 min 100% to 10% solvent B.

The LC system was coupled on-line to an Agilent 6490 QQQ mass spectrometer. Parameters were set as peak width 0.02 min, fragmentor 220V, positive polarity, drying gas 10 L/min, nebulizer 20 psi, drying gas temperature 350°C. Qualitative analysis of neutral lipids is performed by analysis for their unique molecular fragmentation products using a Parent Ion scan of common fragment ions characteristic for the particular class of neutral lipids. Each molecular species is uniquely identified by the parent-to-daughter ion mass transition and for the specific LC retention time.

### Image analysis

Image analysis was performed using ImageJ to quantify the density of SDS-PAGE gel bands in liposome sedimentation assay. GraphPad Prism 10 was used for all statistical analysis.

## ACKNOWLEDGEMENTS

The authors wish to acknowledge the Stony Brook Cancer Center Biological Mass Spectrometry Shared Resource for expert assistance with lipidomics analysis and the Stony Brook Microscopy Imaging Center for expert assistance with fluorescence microscope. We thank the training and support of Samantha Stettnisch and Sabrina Hafeez from the lab of Dr. Benjamin Martin (Stony Brook) for cell imaging. This work was supported by the NIH grant R35GM128666 (M.V.A.), a Sloan Research Fellowship (M.V.A.), a Carol Baldwin award (D.C.), startup funds from the Stony Brook Cancer Center (D.C.), an American Heart Association Fellowship 23PRE1019634 (L.W.), and the Agence Nationale de la Recherche, ANR-21-CE13-0014-LIPDROPER (A.R.T).

