## Supplemental Information for "DDHD2 possesses both lipase and transacylase capacities that remodel triglyceride acyl chains"

### Supplemental methods

#### Protein expression and purification:

Full length human DDHD2, S351A, D541A and H681A were codon-optimized for expression in *Escherichia coli* and cloned into pET28 that contains a N-terminal 6x His tag. The genes encoding human DDHD2 ΔHelix, mouse ATGL 1-319 and full length human CGI58 were codon-optimized for expression in *Escherichia coli* and cloned into ppSUMO, which is a derivative of pET28 that contains a N-terminal 6x His tag followed by a cleavable SUMO tag. Plasmids were transformed into BL21(DE3) RIPL cells in Terrific Broth with 50 µg/mL kanamycin, cultured at 37°C to an OD600 around 1.8, and protein expression was induced with isopropyl β-D-1-thiogalactopyranoside (IPTG) to a final concentration of 0.5 mM at 15°C for 18 h. Cells were harvested by centrifugation at 4000g for 20mins and stored at -80°C until use. Cell pellets were resuspended with buffer containing 50 mM Tris-HCl pH 8, 500 mM NaCl, 5% glycerol, 1 mM tris(2-carboxyethyl)phosphine (TCEP), 1 mM PMSF. After sonication at 85% amplitude for 10 mins, cell lysates were centrifuged at 68,000 x g at 4°C for 1 h. The supernatant was then incubated with 5 mL washed Ni-NTA resin at 4°C for 1 h prior to loading onto a gravity column. The resin was washed with buffer containing 50 mM Tris-HCl pH 8, 500 mM NaCl, 5% glycerol, 1 mM TCEP, 20 mM imidazole followed by an increase of imidazole concentration to 60 mM for second wash, and 300 mM for elution. If needed, the SUMO tag on eluted protein was cleaved by Ulp-1 protease at 4°C overnight. The eluted proteins were further purified by a HiLoad 26/600 Superdex 200 pg column (GE life sciences), analyzed by SDS-PAGE, concentrated using a centrifuge tube depended on protein sizes (30k or 50k MWCO, Pall corporation) to 1-3 mg/mL, flash-frozen and stored at -80°C.

The gene encoding human HSL was cloned into YM-Bac2, which is a modified version of pFastBac Htb (Invitrogen) that contains a N-terminal 6xHis tag followed by a Dual-Strep tag. Plasmid was expressed in Sf9 cells using baculovirus. 1 L of cells were infected with 1 mL of passage 4 baculovirus at 2 million cells/mL at >95% viability and harvested 48 h later. Cell pellets were resuspended with lysis buffer 50 mM HEPES pH 8.0, 500 mM NaCl, 5% glycerol, 0.02% Glyco-dendrimer (GDN), 5 mM β-mercaptoethanol, and protease inhibitors, lysed by sonication, and centrifuged at 81770 x g at 4°C for 1h. The supernatant was then incubated with 5 mL washed Ni-NTA resin at 4°C for 1 h prior to loading onto a gravity column. The resin was washed with buffer containing 50 mM HEPES pH 8.0, 500 mM NaCl, 5% glycerol, 0.02% GDN, 60 mM imidazole and eluted in the same buffer containing 300 mM imidazole. The eluted proteins were further purified by a HiLoad 26/600 Superdex 200 pg column (GE life sciences), analyzed by SDS-PAGE, concentrated using a centrifuge tube (30k MWCO, Pall corporation) to 1-2 mg/mL, flash-frozen and stored at -80°C.

**Supplemental Table 1. Qualitative mass transitions along with collision energy, applied for the MRM experiments of NBD-labeled lipid products analysis.**

Supplemental Table 1. Qualitative LC/MS parameters for lipid analysis on NBD-lipids

| Lipid | Precursor | Mass transition 1 | Mass transition 2 | Collision energy (eV) | Retention time (min) |
|---|---|---|---|---|---|
| NBD-MAG (10:0) | $[M + NH4]^+$:*442.2* | 442.2-407.1 | N/A | 20 | 0.91 |
| NBD-DAG (10:0/10:0) | $[M + NH4]^+$:*596.2* | 596.2-407.1 | 596.2-229.2 | 20 | 3.28 |
| NBD-DAG (10:0/18:1) | $[M + NH4]^+$:*706.5* | 706.5-671.4 | 706.5-407.1 | 20 | 4.76 |
| NBD-TAG (10:0/10:0/18:1) | $[M + NH4]^+$:*860.6* | 860.6-671.4 | 860.6-561.3 | 20 | 9.27 |
| NBD-TAG (10:0/18:1/18:1) | $[M + NH4]^+$:*970.7* | 970.7-671.4 | N/A | 20 | 10.45 |

**Supplemental Table 2. Qualitative mass transitions along with collision energy, applied for the MRM experiments of unlabeled lipid products analysis.**

Supplemental Table 2. Qualitative LC/MS parameters for lipid analysis on unlabeled lipids

| TAG | Precursor | Mass transition | Collision energy (eV) | Retention time (min) |
|---|---|---|---|---|
| 8:0/8:0/8:0 | $[M + NH4]^+$:*488.4* | 488.4-327.3 | 20 | 6.328 |
| 8:0/8:0/18:1 | $[M + NH4]^+$:*626.5* | 626.5-327.3 | 20 | 10.414 |
| 8:0/8:0/21:1 | $[M + NH4]^+$:*668.5* | 668.5-327.3 | 20 | 10.676 |
| 8:0/18:1/18:1 | $[M + NH4]^+$:*764.7* | 764.7-603.5 | 20 | 11.27 |
| 8:0/18:1/21:1 | $[M + NH4]^+$:*806.7* | 806.7-645.6 | 20 | 11.532 |
| 8:0/21:1/21:1 | $[M + NH4]^+$:*848.8* | 848.8-687.6 | 20 | 11.999 |
| 18:1/18:1/18:1 | $[M + NH4]^+$:*902.8* | 902.8-603.5 | 20 | 12.131 |
| 18:1/18:1/21:1 | $[M + NH4]^+$:*944.8* | 944.8-603.5 | 20 | 12.261 |
| 18:1/21:1/21:1 | $[M + NH4]^+$:*986.9* | 986.9-687.6 | 20 | 12.526 |
| 21:1/21:1/21:1 | $[M + NH4]^+$:*1029.1* | 1029.1-687.6 | 20 | 12.657 |
| **DAG** | **Precursor** | **Mass transition** | **Collision energy (eV)** | **Retention time (min)** |
| 8:0/8:0 | $[M + NH4]^+$:*362.3* | 362.3-201.2 | 20 | 3.585 |
| 8:0/18:1 | $[M + NH4]^+$:*500.4* | 500.4-339.3 | 20 | 6.929 |
| 8:0/21:1 | $[M + NH4]^+$:*542.4* | 542.4-381.3 | 20 | 8.94 |
| 18:1/18:1 | $[M + NH4]^+$:*638.6* | 638.6-339.3 | 20 | 10.545 |
| 18:1/21:1 | $[M + NH4]^+$:*680.6* | 680.6-339.3 | 20 | 10.74 |
| 21:1/21:1 | $[M + NH4]^+$:*722.6* | 722.6-381.3 | 20 | 10.938 |

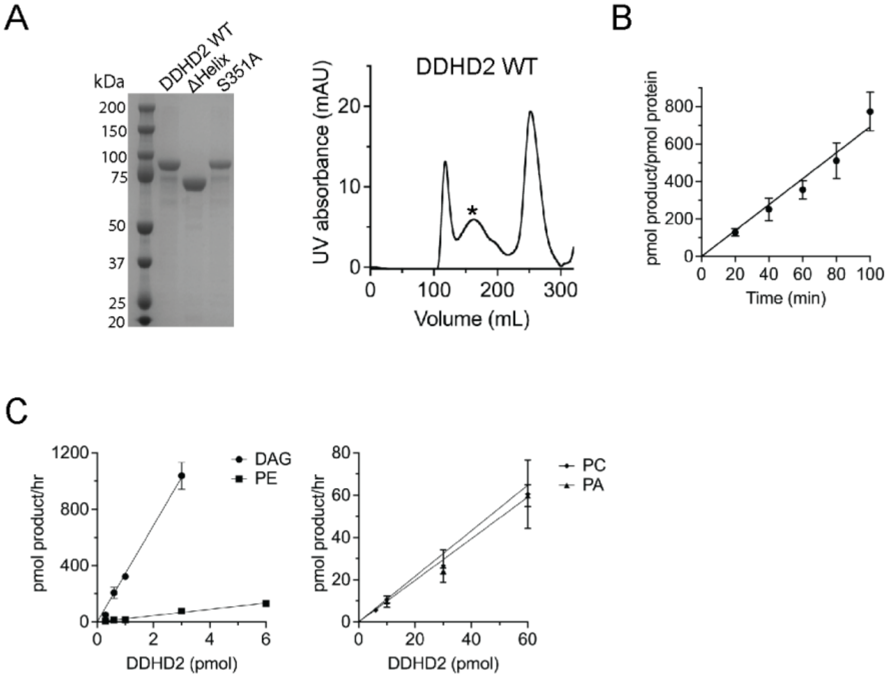


**Supplemental Figure 1.** (A) Left: SDS-PAGE for purified DDHD2 WT, ΔHelix and S351A variants. Right: Representative size exclusion chromatography for DDHD2 WT. Asterisk (*) indicates peak fractions collected. (B) Time linearity of NBD-DAG in Triton X-100 mixed micelles from 0 to 100 minutes.
(C) Linear range of DDHD2 activity towards DAG and PE (left), PA and PC (right). Error bars represent standard deviation (n=3).

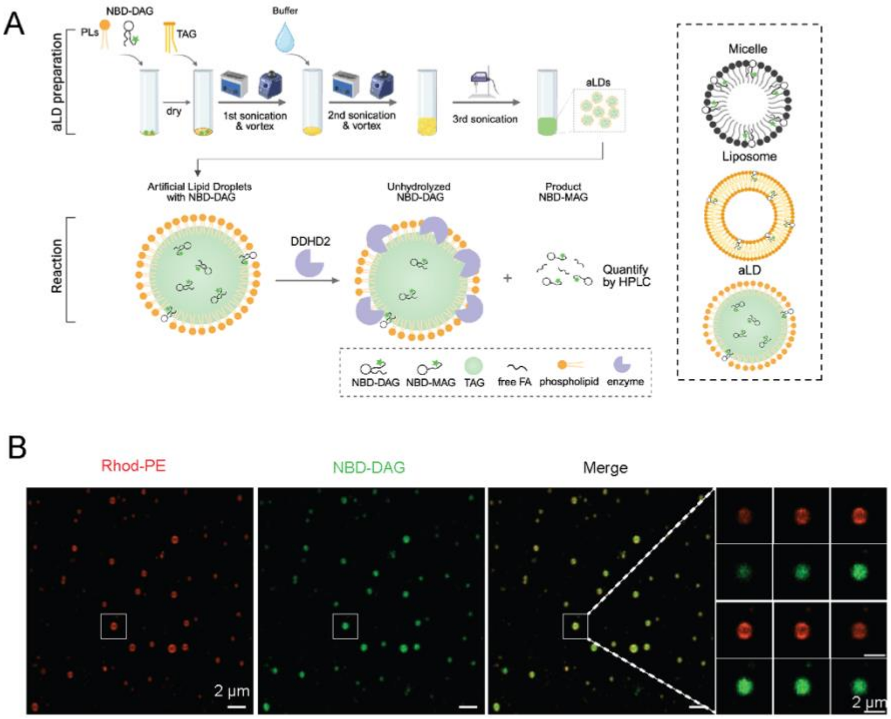


**Supplemental Figure 2.** (A) In vitro lipid droplet generation protocol. (B) Fluorescence imaging of generated lipid droplets using Rhodamine-PE (red) and NBD-DAG (green). Right: Zoom in of one lipid droplet from white box with z-stack from bottom to top. Scale bar: 2 μm.

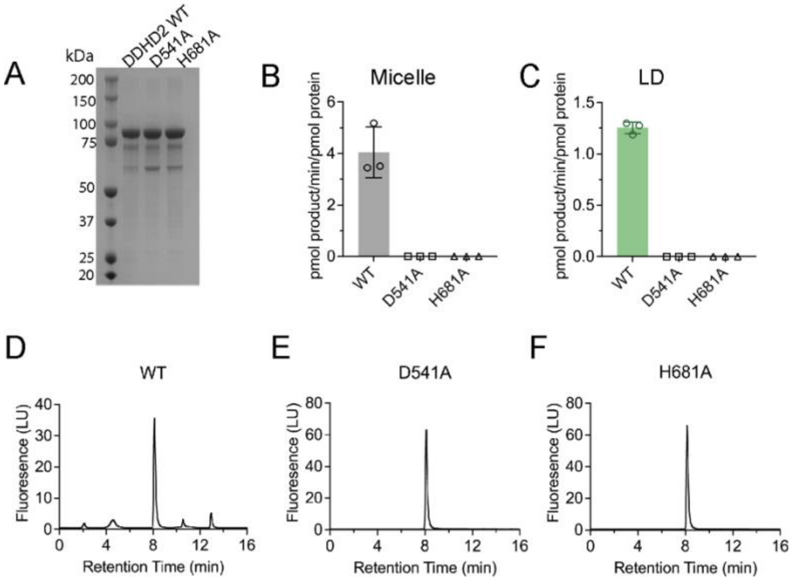


**Supplemental Figure 3.** (A) SDS-PAGE for purified DDHD2 WT, D541A and H681A variants. (B-C) Quantification of DDHD2 variants activity compared to WT in (B)Triton X-100 mixed micelles and (C) artificial lipid droplets. Error bars represent standard deviation (n=3). (D-F) Representative HPLC chromatograms of (D) WT, (E) D541A, and (F) H681A in lipid droplets.

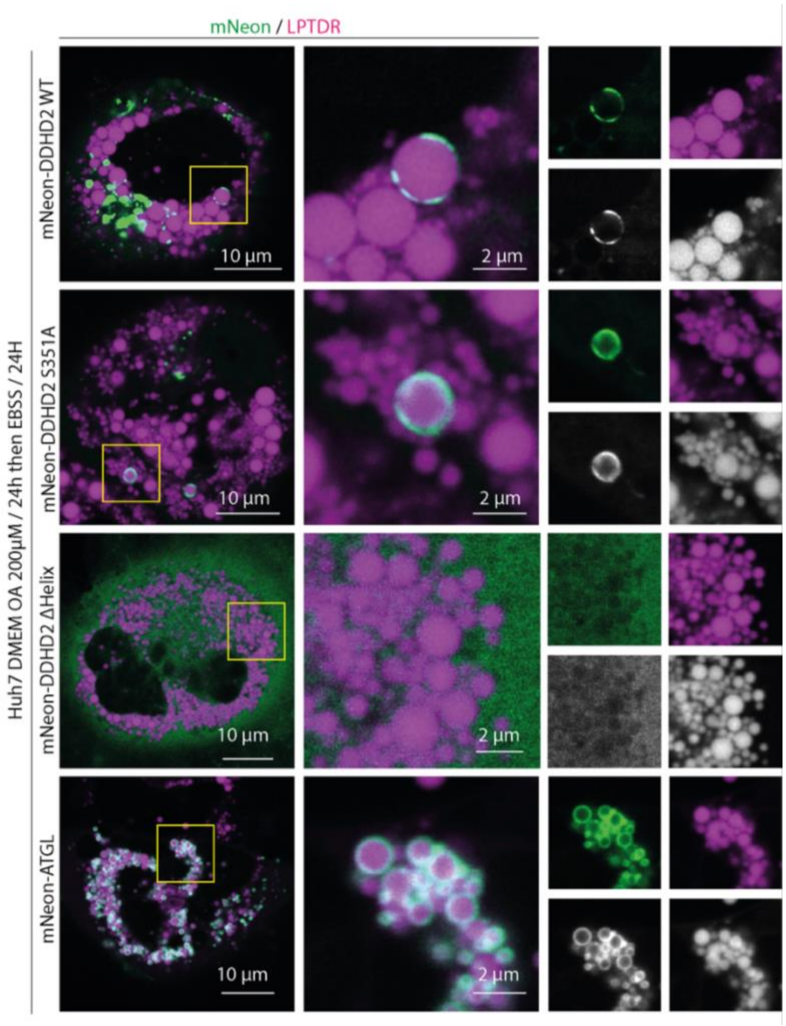


**Supplemental Figure 4.** Representative Airyscan images of Huh7 cells stained by LipidTOX Deep Red (magenta) with overexpression of mNeonGreen-DDHD2 WT, mNeonGreen-DDHD2 S351A, mNeonGreen-DDHD2 ΔHelix and mNeonGreen-ATGL (cyan). Scale bars, 10 µm and 2 µm as labeled.

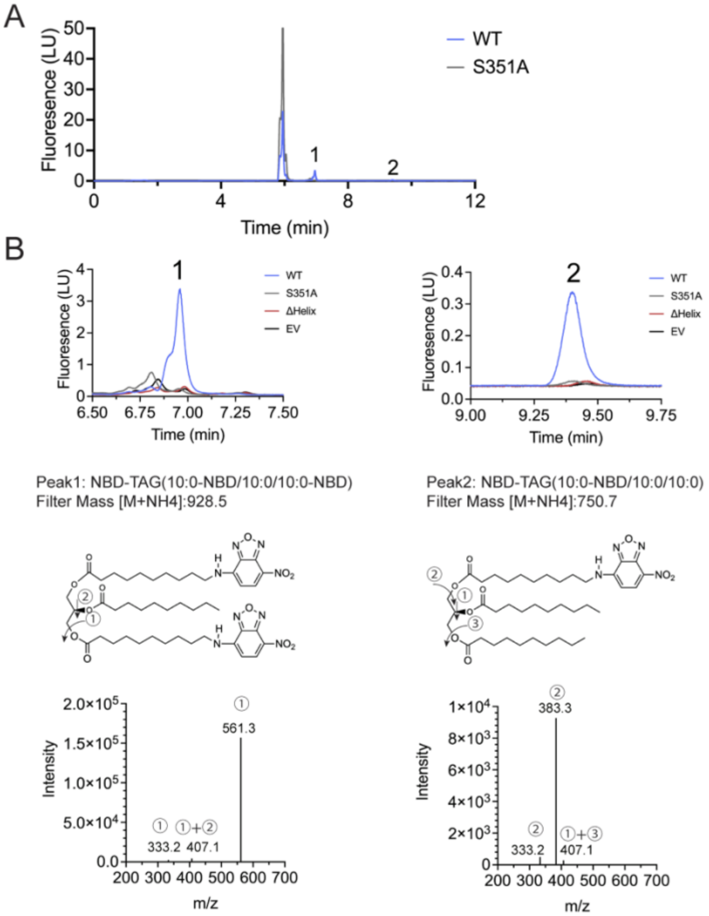


**Supplemental Figure 5.** (A) Representative HPLC fluorescence chromatogram of the cell-based activity assay of DDHD2 WT and inactive mutant S351A in HEK293-T cells. (B) Two peaks generated only in DDHD2 WT are zoomed in. Peak 1 NBD-TAG (10:0-NBD/10:0/10:0-NBD) and peak 2 NBD-TAG (10:0-NBD/10:0/10:0) filtered by mass [M+NH4+] indicated respectively, after collision energy 20 eV applied, molecule breaks down at position 1, 2 or 3, detected mass indicated respectively. These peaks are in absence in S351A, ΔHelix and empty vector samples.
